# Vascular endothelial growth factor A contributes to the increasing of mammalian respiratory epithelial permeability induced by *Pasteurella multocida* infection

**DOI:** 10.1101/2022.10.28.514333

**Authors:** Lin Lin, Jie Yang, Dajun Zhang, Qingjie Lv, Fei Wang, Peng Liu, Mixue Wang, Congcong Shi, Xi Huang, Wan Liang, Chen Tan, Xiangru Wang, Huanchun Chen, Brenda A Wilson, Bin Wu, Zhong Peng

## Abstract

Infections with *Pasteurella multocida* can cause significant zoonotic respiratory problems in both humans and animals. *In vivo* tests in mouse infection models were used to investigate the mechanisms of respiratory epithelial barrier dysfunction during respiratory bacterial infection with these pathogens. Results revealed that *P. multocida* infection significantly increased epithelial permeability and increased expression of vascular endothelial growth factor A (VEGFA) and endothelial nitric oxide synthase (eNOS) in murine tracheae and lungs. In murine lung epithelial cell (MLE-12) models, *P. multocida* infection decreased the expression of tight junctions (ZO-1) and adherens junctions (β-catenin, E-cadherin), but induced the activation of the hypoxia-inducible factor-1α (HIF-1α) and VEGFA signaling. When expression of HIF-1α is suppressed, the induction of VEGFA and ZO-1expression by *P. multocida* infection is decreased. We also found that intervention of HIF-1α and VEGFA signaling affected infection outcomes caused by respiratory bacteria in mouse models. Most importantly, we demonstrated that *P. multocida* infection increased permeability of human respiratory epithelial cells and this process was associated with the activation of the HIF-1α and VEGFA signaling and likely contributes to the pathogenesis of *P. multocida* in humans.

**Importance:** Mammalian respiratory epithelium forms the first line of defense against infections with *Pasteurella multocida*, an important zoonotic respiratory pathogen. In this study, we found *P. multocida* infection increased respiratory epithelial permeability and promoted the induction of the hypoxia-HIF-1α-VEGFA axis in both mouse and murine cell models. Similar findings were also demonstrated in human respiratory epithelial cells. The results from this study gain important knowledge about the pathogenesis of *P. multocida* causing infections in both animals and humans.

## Introduction

Respiratory infections pose serious health and socioeconomic problems owing to their associated high mortality rates and economic costs (1). Respiratory infections are estimated to be responsible for annual deaths of 3.96 million people worldwide (2). As of 19 July 2022, the ongoing COVID-19 pandemic has resulted in 559,469,605 confirmed cases and 6,361,157 confirmed deaths globally, as reported by the World Health Organization (WHO) (3). In agriculture, respiratory disorders such as avian influenza (AI), bovine respiratory disease (BRD), and porcine respiratory disease complex (PRDC) have resulted in hundreds of millions of economic losses in the global livestock industry (4-6).

The mammalian respiratory epithelium forms the first line of defense against the invasion and damage caused by respiratory pathogens (7, 8). Respiratory epithelial cells are connected to their neighbors by cell-to-cell junctions, including tight junctions (TJs), adherens junctions (AJs), gap junctions, and desmosomes, and these interconnections are the central components of the host respiratory epithelial barrier (8). To invade the body and cause disease, it is necessary for respiratory pathogens to induce the permeability of host respiratory epithelial barriers to promote disease pathogenesis (9). However, the detailed mechanisms used by respiratory pathogens to invade host respiratory epithelial barriers remain unclear.

Vascular endothelial growth factor A (VEGFA) is an endothelial-specific growth factor that accelerates angiogenesis and vascular permeability (10, 11), and endothelial nitric oxide synthase (eNOS) plays an essential role in mediating angiogenesis and endothelial function induced by VEGFA, through the production of nitric oxide (NO) (12). The production of VEGFA is regulated by hypoxia-inducible factor (HIF)-1, a major transcription factor composed of two subunits (α and β) that induces the cellular response to hypoxic conditions (10, 13). It has been found that HIF-1-VEGFA signaling is involved in the development of many diseases, including chronic airway inflammatory diseases (13), *Clostridium difficile* induced enteritis (14), and pneumococcal meningitis (15). However, it remains to be further explored whether HIF-1-VEGFA signaling contributes to the modulation of epithelial permeability induced by respiratory bacterial infection.

*Pasteurella multocida* is an important zoonotic respiratory pathogen that is capable of causing respiratory diseases in multiple animals as well as in humans (16). Isolates belonging to this gram-negative bacterial species are commonly divided into five capsular serogroups/genotypes (A, B, D, E, F) (17-19). Generally, *P. multocida* capsular serogroups/genotypes A, D, and F are commonly associated with respiratory syndromes in both animals and humans, while strains of serogroups/genotypes B and E are frequently associated with hemorrhagic septicaemia (19). In this study, we used *P. multocida* as a model to investigate the mechanisms of respiratory epithelial barrier dysfunction during respiratory bacterial infection. Our results demonstrated that *P. multocida* isolates of both human and animal origins could induce the permeability of respiratory epithelial barriers during infection, and this process was associated with the activation of the HIF-1α/VEGFA pathway.

## Results

### Bacterial infection increases the permeability of respiratory epithelial barrier and promotes production of VEGFA in mouse infection models

To explore whether respiratory bacterial infection influences the permeability of mammalian respiratory epithelial barriers, C57BL/6 mice (5∼6-week-old) were intranasally inoculated with *P. multocida* strains from different hosts (HuN001 from human and HN05 from pig), or PBS (negative control) (Fig. 1A). At 48 hours post inoculation (hpi), we assessed the vascular permeability in tracheae and lungs in infected mice by systemic injection of Evans blue, which is an azo dye that has a very high affinity for serum albumin, and has been widely used to assess the change of permeability in physiologic barriers in different disease models (14). The results revealed that the vascular permeability in tracheae and lungs was remarkably increased in mice infected with respiratory bacteria compared to that of the control mice treated with PBS (*P*< 0.001 for both tracheae and lungs, Fig. 1B, C and D). Production of VEGFA in tracheae and lungs of experimental mice, assayed using ELISA, revealed increased VEGFA levels in bacterially infected mice (Fig. 1E). To observe whether the above findings are also present during the other respiratory bacterial infection, a well-known respiratory bacterial pathogen *Streptococcus pneumoniae* was included for comparison (Fig. 1). The results demonstrated that *S. pneumoniae* infection likewise induced an increase in vascular permeability in murine tracheae and lungs as well as increased VEGFA levels (Fig. 1B, C, D, E), suggesting that increasing vascular permeability in main respiratory organs commonly occurred during respiratory bacterial infection. Immunohistochemical (IHC) analysis using von Willebrand Factor (vWF) antibody (an established microvascular endothelial marker indicating angiogenesis in vascular permeability in tracheae and lungs (14)) revealed an increase in angiogenesis in tracheae and lungs of the infected mice compared to those of PBS-treated mice (Fig. 2); IHC staining with eNOS polyclonal antibody also showed an increased production of eNOS in tracheae and lungs of the bacterially infected mice compared to those of PBS-treated mice (Fig. 2). These findings indicated that increased vascular permeability and angiogenesis occurred in tracheae and lungs during respiratory bacterial infection.

**Fig. 1.**
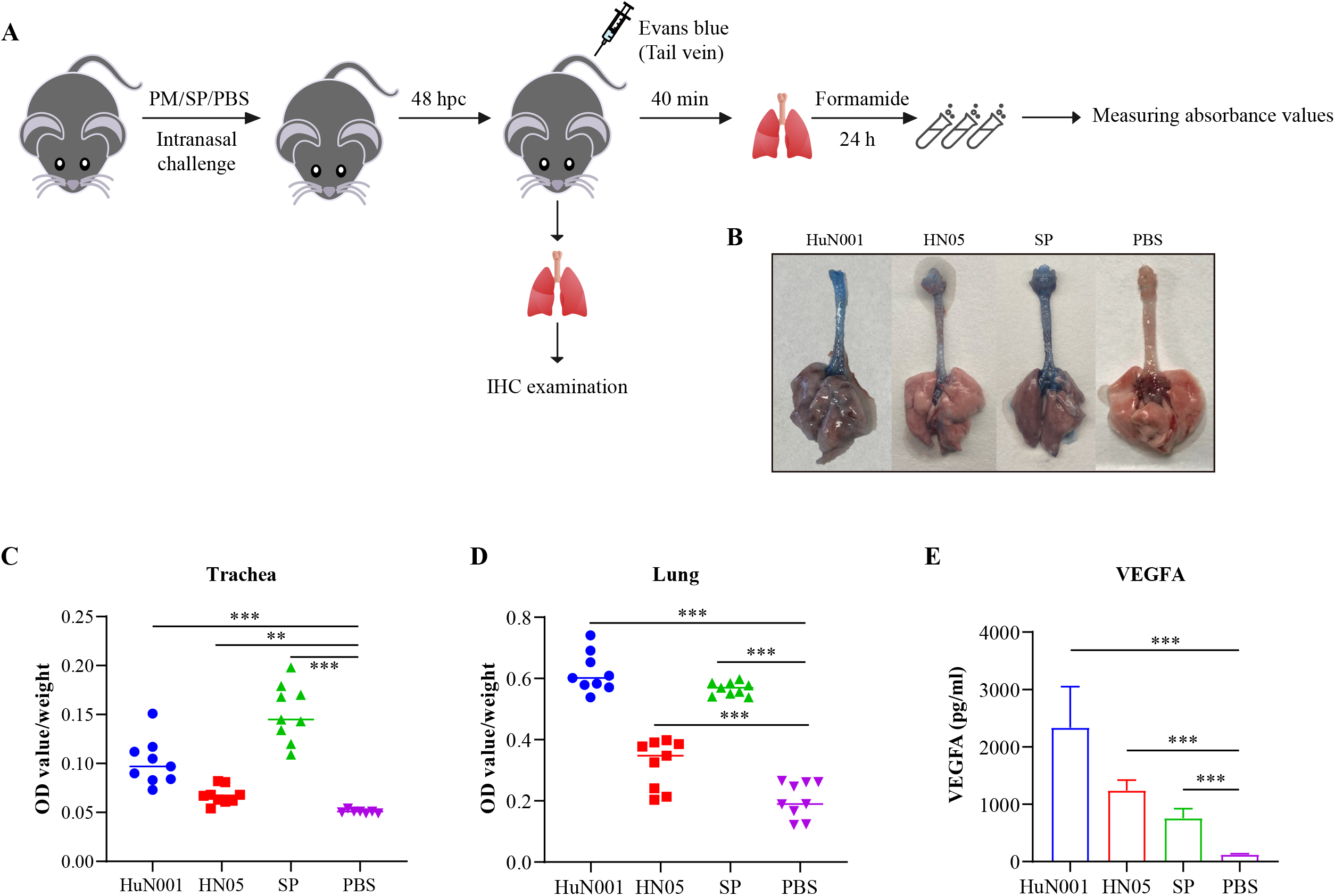
*Pasteurella multocida* (Pm) and *Streptococcus pneumoniae* (Sp) infection increases respiratory vascular permeability in mice. (A) Study design of the mouse experiment. Mice were inoculated intranasally with *P. multocida* HuN001, HN05, *S. pneumoniae* D39, or PBS. After 48 hours, each of the mice were administered Evans blue through the tail vein to assess the vascular permeability in tracheae and lungs. After 40 min, the mice were euthanized and dyes in tracheae and lungs were extracted using formamide. Changes in vascular permeability were evaluated by measuring the absorbance values (OD_620_) of the solutions. In parallel, tracheae and lungs of bacterial infected mice without tail vein injection of Evans blue were also collected for IHC examinations. (B) Visualization of tracheae and lungs obtained from mice inoculated with *P. multocida* strains (HuN001 and HN05) or *S. pneumoniae* (SP), and the control (PBS), showing respiratory vasculature and staining with Evans blue dye. (C) Quantification of Evans blue in tracheae obtained from bacterially infected mice and control mice. (D) Quantification of Evans blue in lungs obtained from bacterially infected mice and control mice. (E) The expression of VEGFA in tracheae and lungs obtained from bacterially infected mice and the control mice. IHC: Immunohistochemical; hpc: hours post challenge; VEGFA: Vascular endothelial growth factor A.

**Fig. 2.**
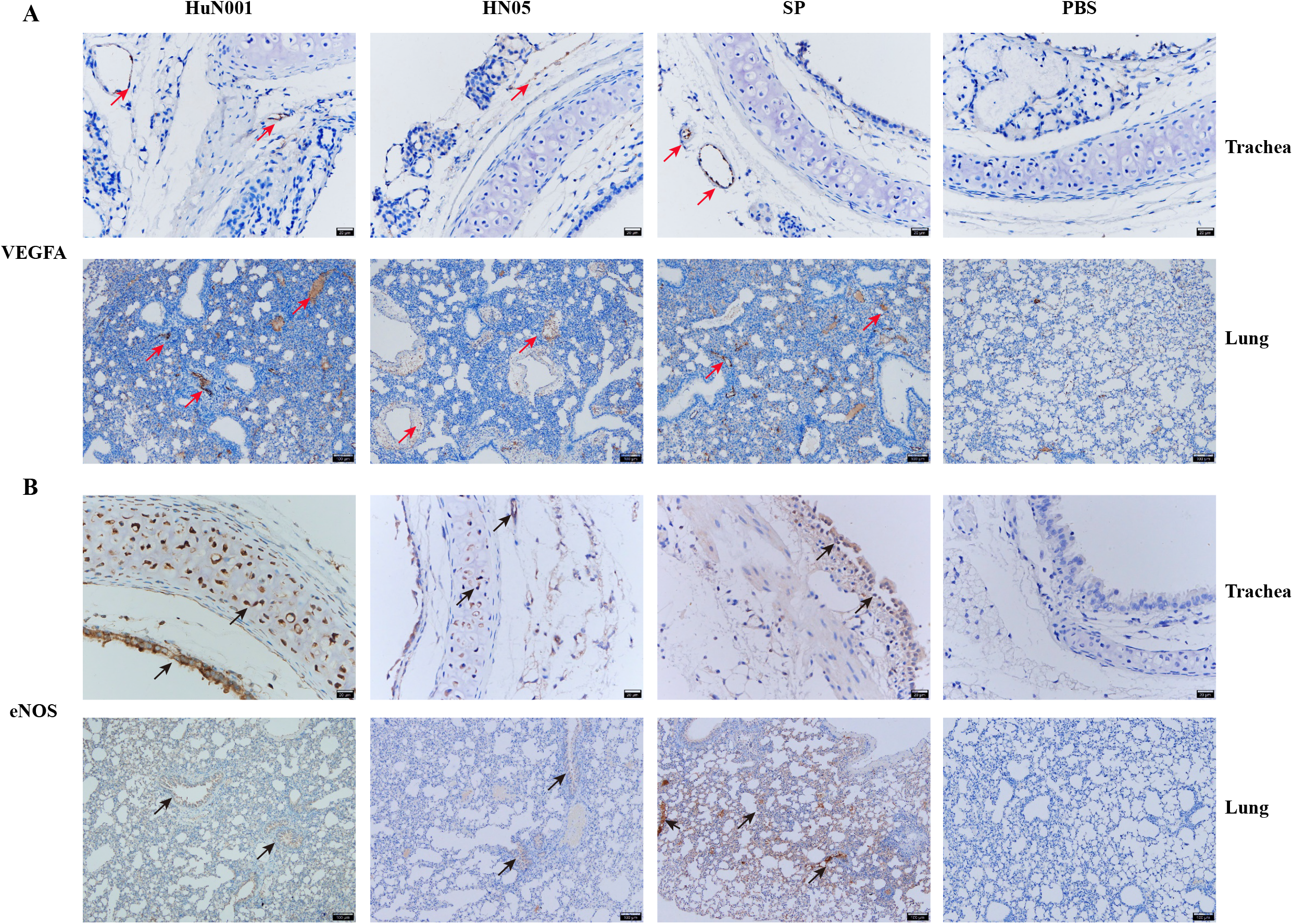
Infection with *Pasteurella multocida* or *Streptococcus pneumoniae* increases VEGFA and eNOS expression in mouse tracheae and lungs. Shown are images from immunohistochemical examination of tracheae and lungs obtained from bacterially infected mice, compared to control mice, treated as described in Figure 1. (A) The expression of VEGFA (shown with the red arrows; marked using the von Willebrand Factor (vWF) antibody) in tracheae (top panel) and lungs (bottom panel) obtained from bacterially infected mice (HuN001, HN05, Sp) and the control mice (PBS); (B) The expression of eNOS (shown with the black arrows; marked using the eNOS polyclonal antibody) in tracheae (top panel) and lungs (bottom panel) obtained from bacterial infected mice and the control mice.

### *P. multocida* infection induces disruption of barrier functions of murine respiratory epithelial cells by promoting the HIF-1α-VEGFA signaling

We next investigated the impact of respiratory bacterial infection on the permeability of mammalian respiratory epithelial barriers. To achieve this, monolayers of murine lung epithelial cells (MLE-12) were infected with *P. multocida* isolates from different hosts or *S. pneumoniae*. After infection for 6 h, expression levels of key TJ proteins (ZO-1) and AJ proteins (β-catenin, E-cadherin) were determined using either qPCR or Western-blotting analysis. The results showed that expression of these proteins in respiratory bacterially infected cells were remarkably decreased compared to those in PBS-treated cells (Fig. 3A and B). Visualization by using laser confocal microscopy also revealed that *P. multocida* infection decreased the expression of the key TJ protein ZO-1 (Fig. 3C).

**Fig. 3.**
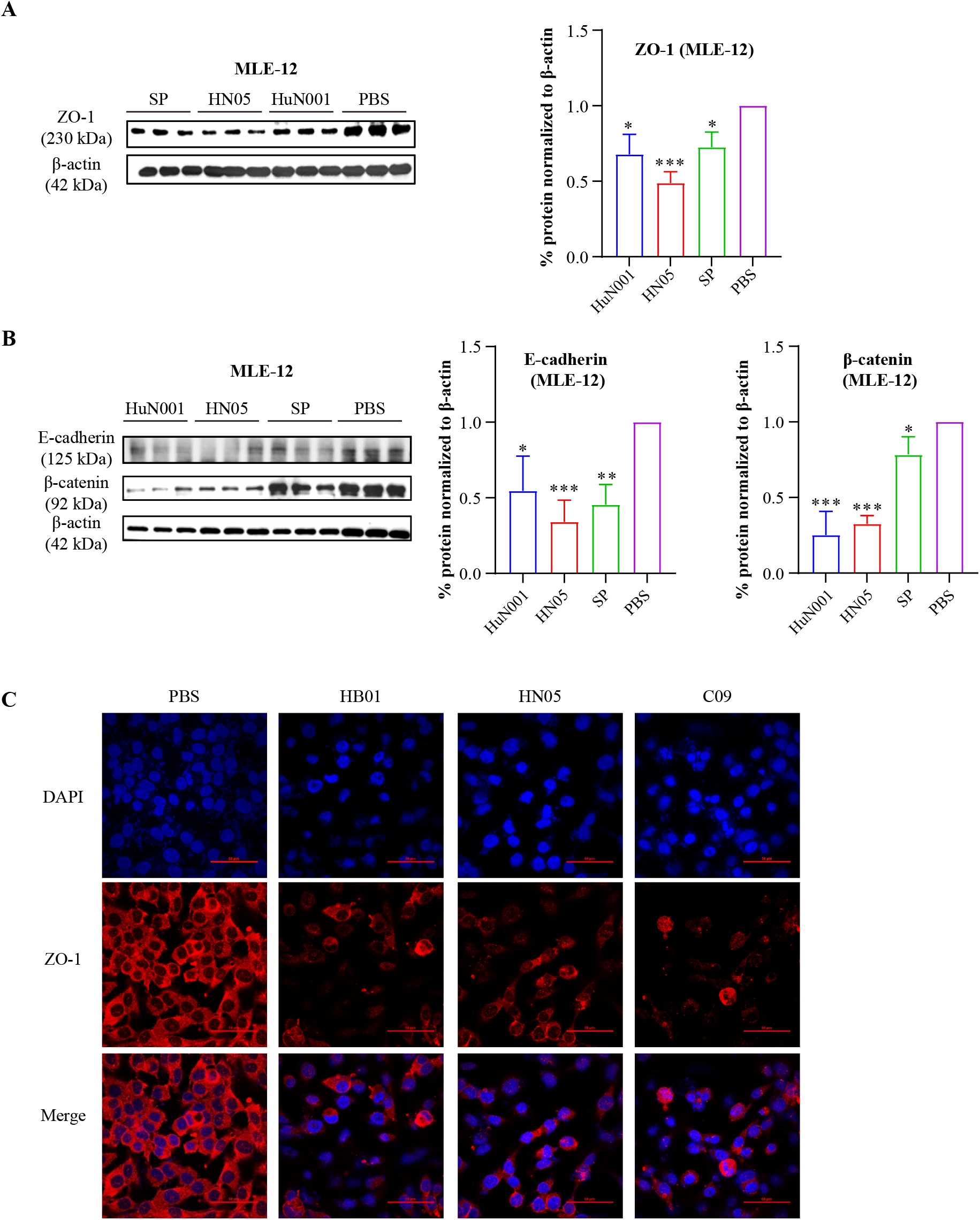
Decreased expression of proteins at tight junctions (ZO-1) and adherens junctions (β-catenin, E-cadherin) in murine lung epithelial cells (MLE-12) infected with *Pasteurella multocida* strains (HuN001, HN05) or *Streptococcus pneumoniae* (Sp). (A) Expression of ZO-1 in bacterially infected MLE-12 cells compared to PBS-treated cells. Right is a plot showing quantifications of the western blot from cell extracts in triplicate. (B) Western blots demonstrating the expression of β-catenin and E-cadherin in bacterially infected cells and PBS-treated cells; (C) Laser confocal microscopy images showing reduced expression of ZO-1 in MLE-12 cells infected with *P. multocida* strains (HB01 from cattle; HN05 from pigs; C09 from cats) compared to control PBS-treated cells. DAPI, 4’,6-diamidino-2-phenylindole.

To explore the involvement of HIF-1α-VEGF signaling pathway in mammalian respiratory barrier dysfunction during respiratory bacterial infection, we examined the expression levels of VEGFA and HIF-1α in MLE-12 cells post *P. multocida* or *S. pneumoniae* infection for 6 h. The results showed that expression of HIF-1α and VEGFA remarkably increased in respiratory bacterially infected murine respiratory epithelial cells compared to those treated with PBS (Fig. 4A, B, C, and D). Next, we knocked-down HIF-1α expression in MLE-12 cells and determined the expression levels of VEGFA in the cells post *P. multocida* infection. The results revealed that the expression of VEGFA decreased in HIF-1α-knock-down cells compared to that in the wildtype cells after bacterial infection (Fig. 4E and F). In contrast, expression of the TJ protein ZO-1 in HIF-1α-knock-down cells did not decrease as much after infection (Fig. 4F). The above findings indicated *P. multocida* infection led to disruption of the barrier functions of murine respiratory epithelial cells, by downregulating the expression of TJ and AJ proteins between the cells; and this process was associated with the activation of the HIF-1α/VEGF pathway induced by the bacterial infection.

**Fig. 4.**
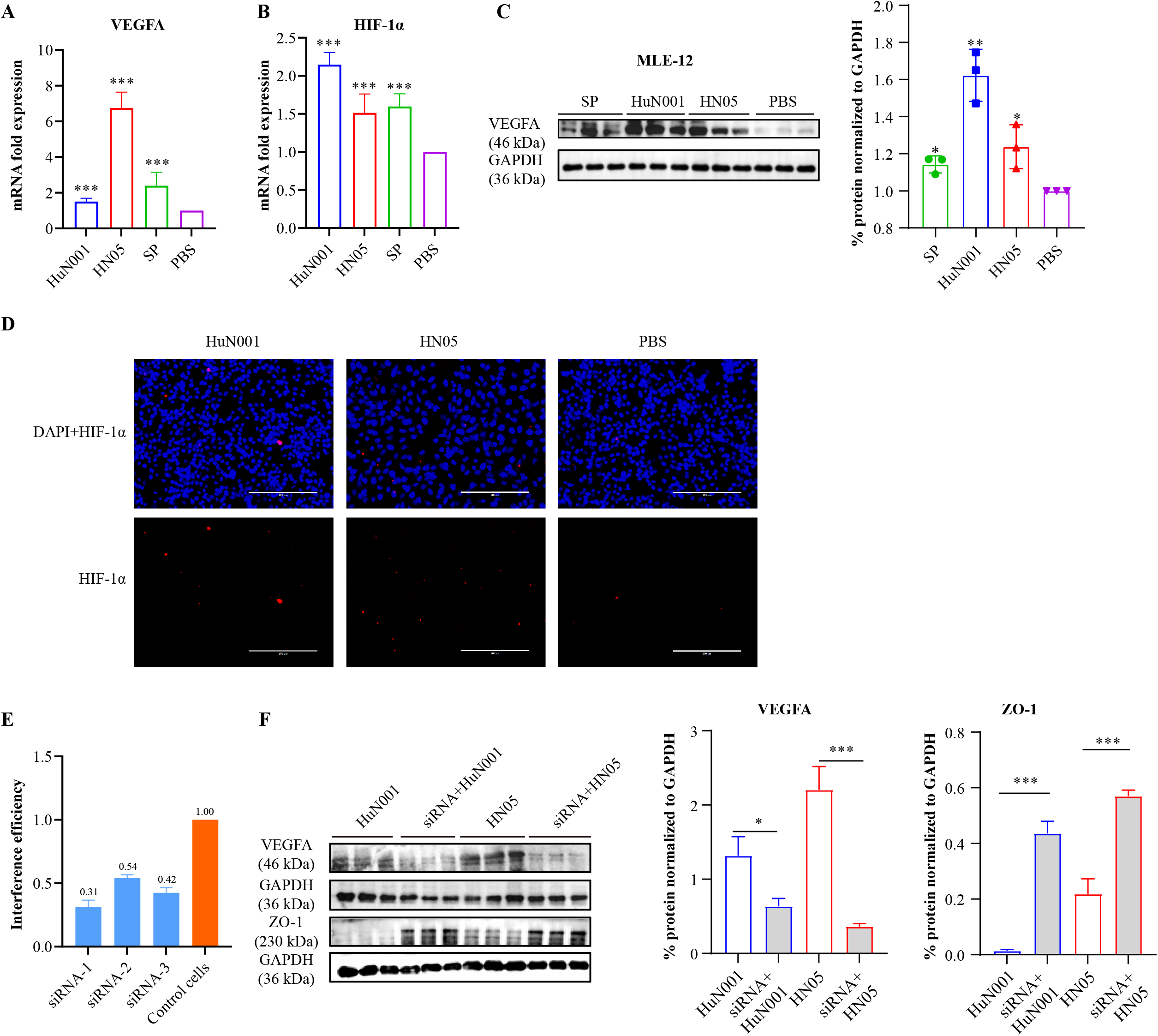
Expression of VEGFA and HIF-1α in murine lung epithelial cells (MLE-12) infected with *Pasteurella multocida* strains (HuN001, HN05) or *Streptococcus pneumoniae* (Sp) for 6 hours. (A) Transcription of VEGFA in bacterially infected cells compared to PBS-treated cells, as determined by qPCR. (B) Transcription of HIF-1α in bacterially infected cells compared to PBS-treated cells, as determined by qPCR. (C) Western blots demonstrating the expression of VEGFA in bacterially infected cells compared to PBS-treated cells. Right is plot showing quantifications of the western blot from cell extracts in triplicate. (D) Increased transcription of HIF-1α in bacterially infected cells compared to PBS-treated cells, as determined by immunofluorescence. (E) Efficacy of three specific siRNAs in suppressing the expression of HIF-1α in MLE-12 cells. (F) Expression of VEGFA and ZO-1 in HIF-1α-suppressed cells after infection with *P. multocida* strains (HuN001, HN05) for 6 hours. Right are plots showing quantifications of the western blot (left panels) from cell extracts in triplicate (control in duplicate).

### Treatment with a VEGFA-blocking antibody attenuates vascular permeability and offers protection against respiratory bacteria in mouse infection models

We next explored whether blocking the HIF-1α-VEGFA signaling pathway could relieve respiratory bacterial infections. To achieve this, C57BL/6 mice (5∼6-week-old; n=10) were first intranasally infected with *P. multocida* strain HN05 (7.5×10^8^ CFU per mouse), and at 4 hours post infection (hpi), 1 day post infection (dpi), and 2 dpi, each of the infected mice received a treatment with the VEGFR-2 inhibitor SU1498 (30 mg/kg), the HIF-1α agonist DMOG (400mg/kg; of), the eNOS inhibitor L-NAME (20mg/kg), or PBS (100 μl) through intraperitoneal injection (Fig. 5A). Mortality recording revealed that treatment of VEGFR-2 inhibitor (SU1498) or eNOS inhibitor (L-NAME) decreased the death rate of mice induced by *P. multocida* infection, while treatment of HIF-1α agonist (DMOG) accelerated the mortality (Fig. 5B and C). We also compared the pathological damages caused by *P. multocida* and *S. pneumoniae* in tracheae and lungs of mice treated by SU1498 to those of mice without treatment. Histological examinations showed that showed that *P. multocida* or *S. pneumoniae* infection induced severe drop of epithelial cells and inflammatory cells infiltration in murine tracheae; and treatment of SU1498 reduced these damages (Fig. 5D). In murine lungs, respiratory bacterial infection caused severe thickening of alveolar walls and fibrosis, as well as exudation of inflammatory cells and red cells in alveolar spaces (Fig. 5E). In contrast, treatment with SU1498 reduced these damages (Fig. 5E). The above findings indicated that the HIF-1α-VEGFA signaling pathway contributed to the pathogenesis of bacterial pathogens causing respiratory infections.

**Fig. 5.**
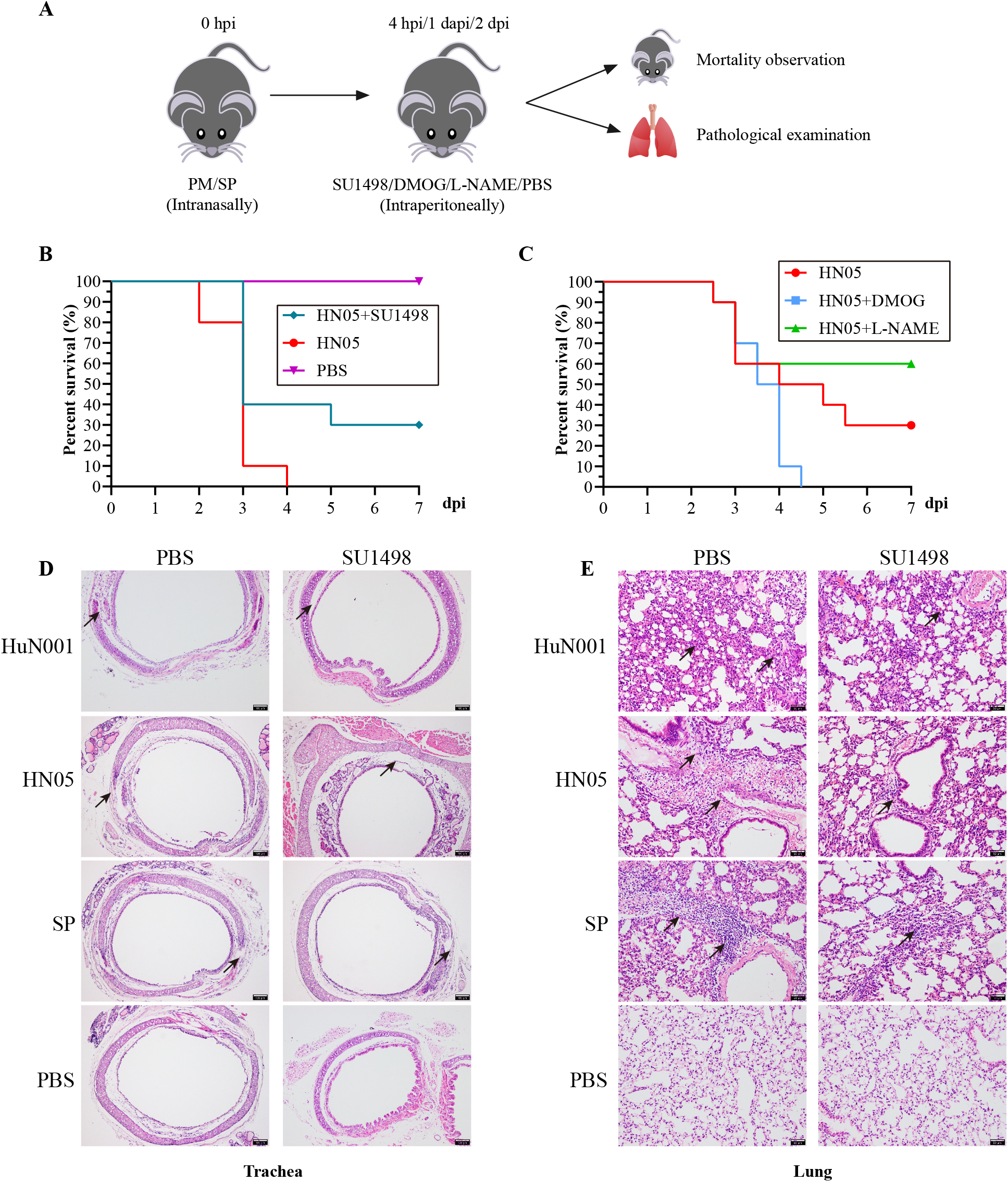
The effect of intervention of HIF-1α/VEGFA signaling on infections caused by respiratory bacteria in mouse models. (A) Study design of the mouse experiments; (B) Survival curves of *P. multocida* infected mice with/without the treatment of the inhibitor of VEGFR-2 (SU1498); (C) Survival curves of *P. multocida* infected mice with/without the treatment of the agonist of HIF-1α (DMOG) or the inhibitor of eNOS (L-NAME); (D) Histological analysis of the tracheae obtained from bacterially infected mice with/without treatment of VEGFR-2 inhibitor (SU1498) (HE 100×); (E) Histological analysis of the lungs obtained from bacterially infected mice with/without the treatment of VEGFR-2 inhibitor (SU1498) (HE 100×); Pm: *Pasteurella multocida*; Sp: *Streptococcus pneumoniae*. Histological damages are shown using black arrows.

### Infections with *P. multocida* isolates from multiple hosts alter barrier functions and activate the HIF-1α-VEGFA pathway in human respiratory epithelial cells

We used human respiratory epithelial cells (BEAS-2B) as a model to tentatively explore the mechanisms of *P. multocida* pathogenesis. Since our recent work demonstrated that *P. multocida* isolates from different hosts are capable of adhering and invading BEAS-2B cells (20), we first examined expression levels of TJ proteins (ZO-1, Occludin) and AJ proteins (β-catenin, E-cadherin) in BEAS-2B cells infected with *P. multocida* isolates from different hosts (HuN001 from humans, C09 from cats, C48-1 from poultry, HB01 from cattle, or HN05 and HN06 from pigs). The results revealed that *P. multocida* from different hosts showed capacities of invading human respiratory epithelial cells, and among the non-human isolates, *P. multocida* from cat displayed the strongest capacity to infect BEAS-2B cells, followed those from poultry, cattle, and pigs, respectively (Fig. 6A). Following this, we also examine the activation of HIF-1α-VEGFA signaling and found that the expression levels of VEGFA and HIF-1α were increased in the cells infected with *P. multocida* from different hosts compared to their expression in the PBS-treated cells (Fig. 6B, C, and D).

**Fig. 6.**
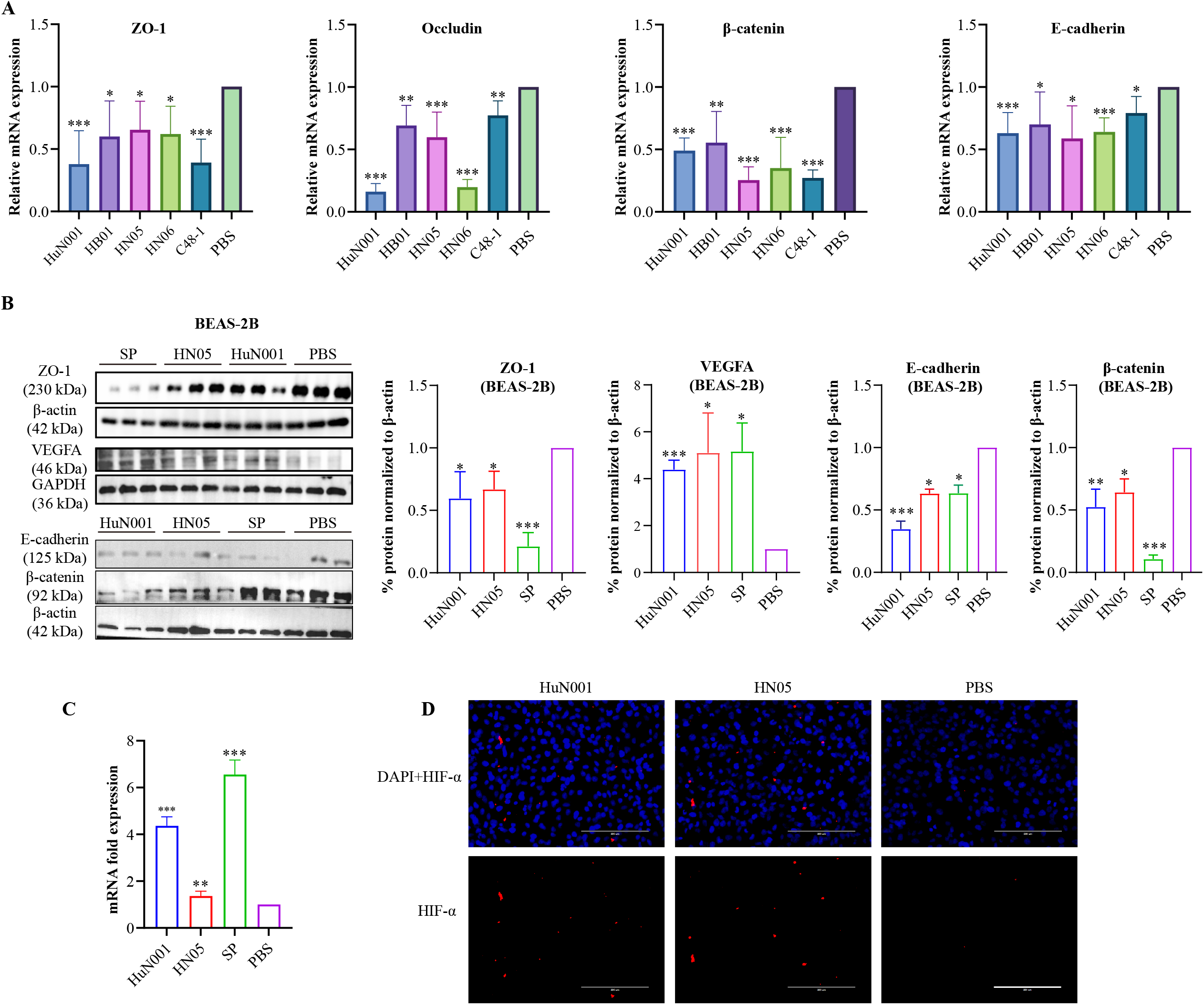
Induction of vascular permeability and VEGFA/HIF-1α signaling in human respiratory epithelial cells (BEAS-2B) infected with *Pasteurella multocida* strains or *Streptococcus pneumoniae*. (A) Transcription of tight junction proteins (ZO-1, occludin) and adherens junction proteins (β-catenin, E-cadherin) in cells infected with *P. multocida* isolates from different hosts (HuN001 from human; HB01 from cattle; HN05 and HN06 from pigs; C48-1 from poultry), or PBS-treated cells, as evidenced by qPCR analysis. (B) Expression of ZO-1, β-catenin, E-cadherin, and VEGFA in bacterially infected cells compared to PBS-treated cells. Right are plots showing quantifications of the western blot (left panels) from cell extracts in triplicate. (C) Transcription of HIF-1α in bacterially infected cells and PBS-treated cells, as determined by qPCR analysis. (D) Immunofluorescence analysis of the transcription of HIF-1α in bacterial infected cells and PBS-treated cells. DAPI: 4’,6-diamidino-2-phenylindole.

## Discussion

As a well-known respiratory pathogen, *P. multocida* is most-commonly infected via respiratory tracts (21). However, the bacterium can cause damages in multiple organs in addition to the respiratory organs, including hearts, spleens, livers, kidneys, guts, and even brains in both humans and animals after infections via respiratory tracts (22-24). In addition, it is common in clinical cases to recover *P. multocida* from organs in addition to the respiratory organs (25-27). Therefore, we hypothesize that *P. multocida* invades and crosses the host respiratory epithelial barrier during infection. However, related studies have not been reported for the pathogenesis of *P. multocida* infection to our knowledge. Our *in vivo* tests demonstrated that *P. multocida* infection, as well as the infection of *S. pneumoniae*, another well-known respiratory pathogenic bacterium associated with respiratory disorders in humans (28), induced an increase in respiratory epithelial permeability in mouse infection models, as evidenced by more Evans blue quantified in the tracheae and lungs of bacterial infected mice (Fig. 1). As an azo dye that has a very high affinity for serum albumin, Evans blue assesses the change of permeability in physiologic barriers such as the blood–brain barrier, intestinal barrier, and respiratory barrier in different disease models (13-15). In addition, we also determined an increased level of VEGFA and eNOS in the tracheae and lungs of mice challenged by both *P. multocida* and *S. pneumoniae* (Fig. 1 and 2). Since protein levels of eNOS are upregulated in response to VEGFA, an increased expression of eNOS indicates an increase in VEGFA expression (12). VEGFA is recognized as a regulatory mediator of vascular permeability, promoting vascular permeability by activating two receptors, VEGFR-1 and VEGFR-2 (29). In many diseases, an increased expression of VEGFA indicates an increased vascular permeability (13-15). Therefore, an increased level of VEGFA and eNOS in the tracheae and lungs of bacterially infected mice also indicates an increase in respiratory epithelial permeability.

Cell-cell junctions between epithelial cells, particularly TJs and AJs, are known to play an essential role in maintaining host epithelial permeability (30, 31). Changes in expression of TJs and AJs between epithelial cell monolayers could be also used to evaluate the condition of epithelial permeability (32, 33). In this study, our *in vitro* tests revealed that *P. multocida* and/or *S. pneumoniae* infection decreased the expression of several key proteins at TJs (ZO-1) and AJs (β-catenin, E-cadherin) in MLE-12 cells (Fig. 3), confirming that respiratory bacterial infection induces the increase of respiratory epithelial permeability. Our *in vitro* tests in MLE-12 cells also demonstrated an increase in VEGFA and HIF-1α in bacterially infected cells (Fig. 4), indicating that the HIF-1α/VEGFA signaling is activated by *P. multocida* and/or *S. pneumoniae* infection. Notably, when the expression of HIF-1α was suppressed, a decreased level of VEGFA and ZO-1 was observed (Fig. 4). As an important signaling pathway involved in perceiving hypoxia, the hypoxia-HIF-1α-VEGF axis has been proposed as an important mechanism driving epithelial barrier disruption and increased permeability of the epithelium (34). This finding has been confirmed in many diseases associated with pathogen infection (14, 15). In agreement with the results of this studies, our findings indicate that respiratory infection induces respiratory epithelial permeability by activating HIF-1α/VEGFA signaling.

Since the above findings suggest an important role of HIF-1α/VEGFA signaling in the pathogenesis of respiratory bacterial infection, we explored whether HIF-1α or VEGFA could be used as targets to relieve respiratory bacterial infection. Our *in vivo* tests in mouse models showed that treatment using an inhibitor of VEGFR-2 (an important receptor of VEGFA) or an inhibitor of eNOS reduced the severity of infection and tissue damage caused by both *P. multocida* and *S. pneumoniae*, while treatment with an agonist of HIF-1α acceleratds the infection and damage (Fig. 5). These findings are similar to those reported from a recent study investigating colonic disease induced by *Clostridium difficile* toxins (14). All these findings suggest that HIF-1α and/or VEGFA represent potential targets to reduce the severity of infections caused by respiratory bacteria and possibly gut pathogens.

While infections caused by *P. multocida* in humans is well documented (16, 35-37), little is known about the mechanisms involved. Our recent work revealed that *P. multocida* isolates from different animals demonstrate capacities of adhering and invading the human respiratory epithelial cells (BEAS-2B) (20). Here using BEAS-2B cells as a model, we demonstrated that infections of *P. multocida* isolates from different animals could induce an increase in respiratory epithelial permeability (Fig. 6), and this process was also associated with the bacterial activation of HIF-1α/VEGFA signaling (Fig. 6).

In summary, our results revealed that respiratory bacterial infection induces activation of HIF-1α/VEGFA signaling that promotes an increase in host respiratory epithelial permeability. Moreover, we demonstrated that *P. multocida* infection increased permeability of human respiratory epithelial cells, which was associated with the activation of HIF-1α/VEGFA signaling, suggesting this may be a mechanism contributing to the pathogenesis of *P. multocida* in humans. We further confirmed that intervention of the HIF-1α/VEGFA signaling could mitigate infections caused by respiratory bacteria in mouse models, thereby presenting this pathway as a suitable candidate for the therapy against respiratory bacterial infections.

## Materials and methods

### Bacterial strains, cell lines, and culture conditions

*P. multocida* strains used in this study are isolated from multiple hosts species: strain HuN001 (GenBank accession no. CP073238) is a capsular genotype A strain recovered from a patient with pneumonia (38); HN05 (GenBank accession no. PPVF00000000) is a capsular genotype D strain recovered from the tracheae of a pig with pneumonia (39); C48-1 is a capsular genotype A strain recovered from fowl cholera; and C09 is a capsular genotype A strain recovered from a cat with respiratory symptoms. *S. pneumoniae* D39 was kindly gifted by Dr. Qi Huang from Huazhong Agricultural University. Unless otherwise specifically notified, *P. multocida* and *S. pneumoniae* strains were cultured on tryptic soy agar (TSA; Becton, Dickinson and Company, MD, USA) or in tryptic soy broth (TSB; Becton, Dickinson and Company, MD, USA) containing 5% bovine serum at 37°C for a minimum of 12 h. MLE-12 cells were maintained in Dulbecco′s Modified Eagle′s Medium/Nutrient Mixture F-12 Ham (DMEM/F-12; Gibco, US) medium containing 2% fetal bovine serum (FBS, Bio-Channel, China), 1% GlutaMAX-1, 1% HEPES 1M buffer solution, 1% penicillin-streptomycin, 1% ITS (Insulin-transferrin-selenium), 10 nM hydrocortisone, and 10 nM estradiol at 37°C under 5% CO_2_ atmosphere; while BEAS-2B cells were maintained in DMEM containing 10% FBS and 1% penicillin-streptomycin at 37°C under 5% CO_2_ atmosphere.

### Animal tests and ethic statements

Mouse experiments were conducted at the Laboratory Animal Center of Huazhong Agricultural University (Wuhan, China) with the approval from the Institutional Ethics Committees (IECs) of the University (approval number: HZAUMO-2021-0142). Laboratory animals were treated following the Regulations on the Administration of Laboratory Animals in Hubei Province [2005]. Briefly, each of the experimental mice (C57BL/6; 5∼6-week-old) were intranasally challenged with *P. multocida* HuN001 (15 CFU), HN05 (5×10^7^ CFU), *S. pneumoniae* D39 (5×10^7^ CFU), or PBS (100 μl). To assess the vascular permeability in tracheae and lungs in challenged mice, each mouse was challenged with Evans blue (30 mg/kg; Sigma, US) through the tail vein at 48 hours post challenge (hpc). After 40 min, the mice were euthanized and dyes in tracheae and lungs were extracted using formamide (2 ml) at 55°C for 24 h (Fig. 1A). Changes in vascular permeability were evaluated by measuring the absorbance values (OD_620_) of the solutions, as described previously (14). In parallel, tracheae and lungs of bacterially infected mice without tail vein injection of Evans blue were also collected for immunohistochemical examinations (Fig. 1A). To investigate the effect of blocking or promotion of the HIF-1α/VEGFA pathway on respiratory bacterial infections in mice, mice infected with HN05 received a treatment of SU1498 (30 mg/kg; inhibitor of VEGFR-2; Catalog NO. 168835-82-3, AdooQ BioScience, US), DMOG (400mg/kg; agonist of HIF-1α; Catalog NO. A13998, AdooQ BioScience, US), L-NAME (20mg/kg; inhibitor of eNOS; Catalog NO. GA11233, GlpBio, US), or PBS (100 μl) through intraperitoneal injection at 4 hpi, 1 dpi, and 2 dpi, respectively (Fig. 5A). Mortalities were recorded every day for seven days post bacterial infection. To evaluate the therapeutic effect after a treatment of SU1498, each of the mice infected by HuN001, HN05, or D39, as well as the PBS-treated mice were euthanized, and their tracheae and lungs were collected for histological examination.

### Histological and Immunohistochemical examinations

Murine tracheae and lungs were fixed using 4% buffered paraformaldehyde for 2 days, and routinely processed for embedding in paraffin and sectioning at 3-5 μm. For histological examination, tissue sections were dewaxed by step-treatments using xylene I (20 min), followed by the treatment of xylene II (20 min), xylene III (20 min), anhydrous ethanol I (5 min), anhydrous ethanol II (5 min), 95% alcohol (5 min), 90% alcohol (5 min), 80% alcohol (5 min), and 70% alcohol (5 min), respectively. After soaking in distilled water for 5 min, the sections were stained using with Harris hematoxylin for 2 min, differentiated with alcohol hydrochloride for 1 s, and washed using water. Thereafter, the sections were stained with 1% water-soluble eosin solution for 3 min, and then washed using water for 30 s. Following dehydration by step treatments using anhydrous ethanol I (5 min), anhydrous ethanol II (5 min), N-butanol (5 min), xylene I (5 min), and xylene II (5 min), the sections were air-dried and sealed using neutral gum for examination under microscope. For immunohistochemical examination, dewaxed sections were dehydrated and heated in a microwave oven in citrate buffer for 15 min for antigen retrieval. After blocking in 3% hydrogen peroxide for 15 min, the sections were overlaid with 10% normal goat serum for 30 min. Following this, the sections were incubated in von Willebrand Factor (vWF) antibody (1:100) (Abcam, UK) or eNOS polyclonal antibody (1:100) (Invitrogen, USA) at 4°C for overnight. The sections were then incubated with HRP-goat-anti-rabbit IgG (diluted 1:300) for 30 min at 37°C and then incubated with 3,3-N-diaminobenzidine tertrahydrochloride (DAB). Finally, the sections were stained with Harris hematoxylin, dehydrated, and sealed for microscope examination.

### Quantitative real-time PCR (qPCR)

Monolayers of MLE-12 and/or BEAS-2B cells were infected with *P. multocida* HuN001 or HN05 (200 MOI) and incubated at 37°C for 6 h. Cells treated with *S. pneumoniae* D39 (OD_600_ = 0.6) or PBS were used as controls. After that, cells were washed with PBS and total RNAs were extracted using the Trizol reagent protocol (Invitrogen, ThermoFisher, Waltham, MA, USA). cDNAs synthesized from the extracted total RNAs using a PrimeScript™ RT reagent Kit with gDNA Eraser (TAKARA, Japan) were used as templates to perform qPCR assays for the checking of the transcriptions of VEGFA, HIF-1α, ZO-1, β-catenin, E-cadherin, and occludin. GAPDH was used as reference gene. Primers used for qPCR are listed in Table S1 in the supplemental material. All experiments were repeated three times.

### Western blotting

Bacterial infected monolayers of MLE-12 and/or BEAS-2B cells were lysed using Radioimmunoprecipitation assay (RIPA) buffer (Beyotime, China) containing protease inhibitors. The products were then centrifuged at 4°C, 12,000 rpm for 10 min, and the concentrations of proteins in the supernatants were measured using a BCA Protein Assay Kit (Beyotime, China). After that, the proteins were separated on a 10% sodium dodecyl sulfate polyacrylamide gel electrophoresis (SDS-PAGE) and then transferred onto a polyvinylidene difluoride (PVDF) membrane (Bio-Rad, USA). The blots were blocked in 5% BSA in Tris-buffered saline with Tween 20 (TBST) for 3 h at room temperature. Following this, the blots were incubated overnight at 4°C with either VEGFA polyclonal antibody (1:2000) (Catalog NO. 19003-1-AP, Proteintech, China), ZO-1 polyclonal antibody (1:2000) (Catalog NO. 21773-1-AP, Proteintech, China), E-cadherin monoclonal antibody (1:1000) (Catalog NO. P12830, Cell Signaling, US), β-catenin polyclonal antibody (1:5000) (Catalog NO. 21773-1-AP, Proteintech, China), HIF-1α antibody (1:200) (Catalog NO. NB100-134, Novus Biologicals, USA), β-actin monoclonal antibody (1:5000) (Catalog NO. 66009-1-lg, Proteintech, China), or GAPDH monoclonal antibody (1:20000) (Catalog NO. 60004-1-lg, Proteintech, China). After washing, the blots were incubated with species-specific horseradish peroxidase-conjugated antibodies and visualized with enhanced chemiluminescence (ECL) reagents (Beyotime, China). All Western blots were quantified using ImageJ software, and the results were analyzed as the relative immunoreactivity of each protein normalized to the respective loading control.

### siRNA Transfection and bacterial infection

Three specific siRNAs against HIF-1α (Table S1 in the supplemental material) were used to suppress the expression of HIF-1α in MLE-12 cells, and according to their efficacy (Fig. 4E), one of the specific siRNAs (forward: 5′-CCA UGU GAC CAU GAG GAA ATT-3′; reverse: 5′-UUU CCU CAU GGU CAC AUG GAT-3′) was selected for the suppression. This siRNA (20 nM) was transfected in MLE-12 cells using Lipofectamine™ 2,000 reagent (Invitrogen, US) following the manufacturer instructions. Scrambled RNA at the same concentration was also transfected as a control. Afterwards, monolayers of both siRNA-transfected cells and the control cells were infected with *P. multocida* HuN001 and HN05 (200 MOI) for 5 h. The cells were lysed using protease inhibitors in RIPA buffer (Beyotime, China). The expression of VEGFA and ZO-1 in bacterial infected cells were determined using western blotting as described above. This experiment was repeated three times.

### Immunofluorescence

To observe the expression of HIF-1α in the cells during bacterial infection, monolayers of MLE-12 and/or BEAS-2B cells were incubated with *P. multocida* HuN001 or HN05 (200 MOI) for 5 h, followed by washing using precooled PBS for 3 times to remove the free bacteria. Bacterial infected cells were then fixed using precooled formaldehyde for 2 h and were blocked in 5% BSA for 2 h at room temperature. Thereafter, the cells were incubated with HIF-1α antibody (1:200) overnight at 4°C. After washing, the cells were incubated with Cy3-labeled goat anti-rabbit IgG (Catalog NO. AS007, ABclonal, US) at 37°C for 30 min in dark place. Finally, the cells were incubated with antifade mounting medium containing DAPI (Beyotime, China) for 30 min at room temperature at dark place and the expression of HIF-1α was observed under an inverted fluorescence microscope.

### Laser confocal microscopy examination

To observe the expression of ZO-1 in cells during bacterial infection, monolayers of MLE-12 on confocal dishes were infected with *P. multocida* HuN001 or HN05 (200 MOI) for 5 h, followed by washing using precooled PBS for 3 times to remove the free bacteria. Bacterial infected cells were then fixed using precooled formaldehyde for 2 h and were blocked in 5% BSA for 2 h at room temperature. Thereafter, the cells were incubated with ZO-1 polyclonal antibody (1:2000) overnight at 4°C. After washing, the cells were incubated with Cy3-labeled goat anti-rabbit IgG at 37°C for 30 min in the dark. Finally, the cells were incubated with antifade mounting medium containing DAPI (Beyotime, China) for 30 min at room temperature in the dark, and the expression of ZO-1 was observed under a super resolution microscope-confocal laser microscope system. The obtained photos were analyzed using a NIS-Elements Viewer 4.20 software (Nikon, Tokyo, Japan).

### Statistical analysis

Statistical analysis was performed through the “Multiple *t*-tests” strategy in GraphPad Prism 8.0 (GraphPad Software, San Diego, CA). Data represents mean ± SD. The significance level was set at *P* < 0.05 (*), *P* < 0.01 (**), or *P* < 0.001 (***).

## Supplementary material

**Table S1 Oligonucleotides used in this study**.

## Acknowledgments

We would like to thank the State Key Laboratory of Agricultural Microbiology Core Facility for assistance in immunofluorescence and laser confocal microscopy examination. We thank Dr. Qi Huang (Huazhong Agricultural University) for the gift of *Streptococcus pneumoniae* D39. This study was supported in part by National Natural Science Foundation (grant no. 31902241), China Postdoctoral Science Foundation (grant no. 2020T130232), Hubei Provincial Key Research and Development Project (grant no. 2021BBA085), and Huazhong Agricultural University (HZAU) startup funding.

## Conflict of interest statement

Wan Liang is currently an employee of Hubei Jin Xu Agricultural Development Limited by Share Ltd., Wuhan, China. The other authors declare that they have no known competing financial interests or personal relationships that could have appeared to influence the work reported in this paper.

